# Atomoxetine modulates the contribution of high- and low-level signals during free viewing of natural images in rhesus monkeys

**DOI:** 10.1101/2020.07.12.195933

**Authors:** Amélie J. Reynaud, Elvio Blini, Eric Koun, Emiliano Macaluso, Martine Meunier, Fadila Hadj-Bouziane

**Author notes:** Corresponding authors: Amelie Reynaud, and Fadila Hadj-Bouziane, - INSERM U1028, CNRS UMR5292, Lyon Neuroscience Research Center, ImpAct Team, 16 avenue Doyen Lépine 69500 BRON, France.

## Abstract

Visuo-spatial attentional orienting is fundamental to selectively process behaviorally relevant information, depending on both low-level visual attributes of stimuli in the environment and higher-level factors, such as goals, expectations and prior knowledge. Growing evidence suggests an impact of the locus-coeruleus-norepinephrine (LC-NE) system in attentional orienting that depends on task-context. Nonetheless, most of previous studies used visual displays encompassing a target and various distractors, often preceded by cues to orient the attentional focus. This emphasizes the contribution of goal-driven processes, at the expense of other factors related to the stimulus content. Here, we aimed to determine the impact of NE on attentional orienting in more naturalistic conditions, using complex images and without any explicit task manipulation. We tested the effects of atomoxetine (ATX) injections, a NE reuptake inhibitor, on four monkeys during free viewing of images belonging to three categories: landscapes, monkey faces and scrambled images. Analyses of the gaze exploration patterns revealed, first, that the monkeys spent more time on each fixation under ATX compared to the control condition, regardless of the image content. Second, we found that, depending on the image content, ATX modulated the impact of low-level visual salience on attentional orienting. This effect correlated with the effect of ATX on the number and duration of fixations. Taken together, our results demonstrate that ATX adjusts the contribution of salience on attentional orienting depending on the image content, indicative of its role in balancing the role of stimulus-driven and top-down control during free viewing of complex stimuli.

## 1. Introduction

When exploring the environment, the brain receives a multitude of information of different modalities. In the visual modality, visuo-spatial attentional orienting is fundamental to selectively process information, depending on the visual attributes of the elements in the environment and our goals and needs. Growing evidence suggest an involvement of the locus-cœruleus-norepinephrine (LC-NE) system in attentional orienting (Clark et al., 1989; Coull et al., 2001; Dragone et al., 2018; Reynaud et al., 2019). Using pharmacological agents, these studies showed that increasing NE transmission improves attentional orienting when the context is predictive, i.e. when a cue accurately predicts the location of the upcoming target in the large majority (80%) of the trials (Clark et al., 1989; Coull et al., 2001; Reynaud et al., 2019). Another recent study also reported larger increase of pupil diameter, often considered as a proxy of the LC-NE activity, in predictive contexts, in which the cue accurately predicted the location of the upcoming target in 80% of the trials, as compared to non-predictive contexts (50%, chance level, (Dragone et al., 2018)). Taken together, these studies are in favor of a role of NE on visuo-spatial attentional orienting that depends on the context in which the task is performed. These studies typically used spatial cueing tasks, in which the focus of attention was explicitly manipulated using spatial cues predicting the location of the upcoming target. In addition, most of these studies used simple visual displays, encompassing one target and potentially a distractor, adapted for laboratory testing. Such settings emphasize the contribution of goal-driven processes limiting the potential contribution of low-level saliency-driven processes, as well as that of other high-level signals related for example to prior knowledge about scene configuration (Henderson, 2017; Oliva and Torralba, 2007).

Here, we aimed to determine the impact of NE in more naturalistic conditions, without any explicit manipulation of the focus of attention to test for a potential effect of this neuromodulator on both high-level and saliency-driven attention control. We tested four monkeys during free viewing of naturalistic images under two conditions: after saline administration, used as a control condition, and after administration of atomoxetine (ATX), a NE re-uptake inhibitor that enhances the level of NE in the brain. The animals were presented with static images that they could freely explore for three seconds, while we measured their gaze position. In order to manipulate the high-level context, we presented animals with three different categories of images: intact images, i.e. landscape and monkey face images, and scrambled landscape images. The scrambled images, which remove the images meaningful information while preserving their low-level visual features, were introduced to allow us to disentangle the contribution of low-and high-level features in the allocation of attention.

We modeled the influence of low-level signals through salience maps that integrate multiple physical characteristics of the visual images (e.g. color, luminance) (Itti et al., 1998). Previous studies have unveiled the contribution of salience maps in the spatiotemporal deployment of attention in natural scenes in humans (Berg et al., 2009; Parkhurst et al., 2002a) and monkeys (Berg et al., 2009; Berger et al., 2012). Low-level and high-level signals correspond respectively to bottom-up and top-down influences on attention control (Buschman and Miller, 2007; Corbetta et al., 2008; Corbetta and Shulman, 2002). It is thought that the integration of these different types of information into priority maps guides the allocation of attention (Bisley and Mirpour, 2019; Itti and Koch, 2000). To the best of our knowledge, no study has explored the contribution of NE onto these types of influences on visuospatial attentional orienting during free viewing of naturalistic images.

Here, we investigated the effect of ATX on the total free exploration duration as well as the number and mean duration of fixations. To assess the degree to which the animals’ gaze was influenced by salient features in the images such as color, intensity and orientation, i.e. to estimate bottom-up influences, we computed the saliency map for each image using the Graph-Based Visual Saliency model (GBVS) (Harel et al., 2006). We then compared the mean saliency of the locations the animals explored in saline versus ATX conditions. Based on the previous findings about the context-dependent role of NE on attentional orienting discussed above, we hypothesized that ATX would influence the way the animals orient their attention depending on the image content. Given the influences of the LC-NE system on sensory regions (Navarra and Waterhouse, 2019; Waterhouse and Navarra, 2019), including the visual cortex, and on frontal regions controlling top-down processes (Arnsten et al., 2012; Berridge and Spencer, 2016), we further postulated that ATX could influence priority maps during free exploration, hence affecting both saliency-driven and top-down spatial orienting.

## 2. Methods

### 2.1. Subjects

Four female rhesus monkeys (*Macaca mulatta*) aged 5-14 years participated to this study (monkeys CA, GU, GE and CE). Animals had free access to water and were maintained on a food regulation schedule, individually optimized to maintain stable motivation across days. This study was conducted in strict accordance with Directive 2010/63/UE of the European Parliament and the Council of 22 September 2010 on the protection of animals used for scientific purposes and approved by the local Committee on the Ethics of Experiments in Animals (C2EA 42 CELYNE).

### 2.2. Experimental set-up

Monkeys were seated in a primate chair in a sphinx position, with the head immobilized via a surgically implanted plastic MRI-compatible head post (CE and GE) or a non-invasive head restraint helmet (CA and GU) (Hadj-Bouziane et al., 2014), in front of a computer screen (distance: 57cm). Eye position was sampled at 120 Hz using an infrared pupil tracking system (ISCAN Raw Eye Movement Data Acquisition Software) interfaced with a program for stimulus delivery and experimental control (Presentation®).

### 2.3. Free Viewing protocol

Prior to the free viewing protocol, we used a 5-point procedure to calibrate the eye-tracker: the central point was at the center of the monitor, and the four other points were presented at 10° eccentricity on the right, left, top and bottom from the central point. During the free viewing protocol, monkeys were first required to fixate a central cross for 500ms to initiate the trial (4° window for CA and GU; 3° window for GE and CE). Then, one image was presented for 3000ms and the monkeys were free to explore it. During the inter-trial interval (800 to 1400ms), the monkeys received a reward regardless of their exploratory pattern during the previous trial. Each testing session comprised the presentation of 30 images, subdivided into 3 categories: 10 monkey face images (584 x 584 pixels, 10° x 10°), 10 natural landscape images (876 x 584 pixels, 15° x 10°), and 10 scrambled landscape images (876 x 584 pixels, 15° x 10°). Each category comprised 5 original images and 5 horizontally flipped images, in order to control for possible biases in the lateralized distribution of objects or salient features across the images (see figure 1A for an example of images presented in one testing session). The monkey face images were the same across all sessions, whereas new landscape and scrambled images were presented at every session. Within each session, the order of the stimuli (and categories) presentation was randomized. Monkey face images were collected from the internet (criteria: rhesus macaque face with a neutral emotion) and the landscape images were drawn from the MIT places database (from 2 categories: badland and cabin outdoor) (Zhou et al., 2017, 2014). The scrambled landscape images were generated by taking the two-dimensional Fourier transform of the natural landscape images, scrambling the phase, and then taking the inverse Fourier transform. This preserves the second order statistics of the images, while interfering with higher-order statistics and - most importantly - making the image content undecipherable.

**Figure 1.**
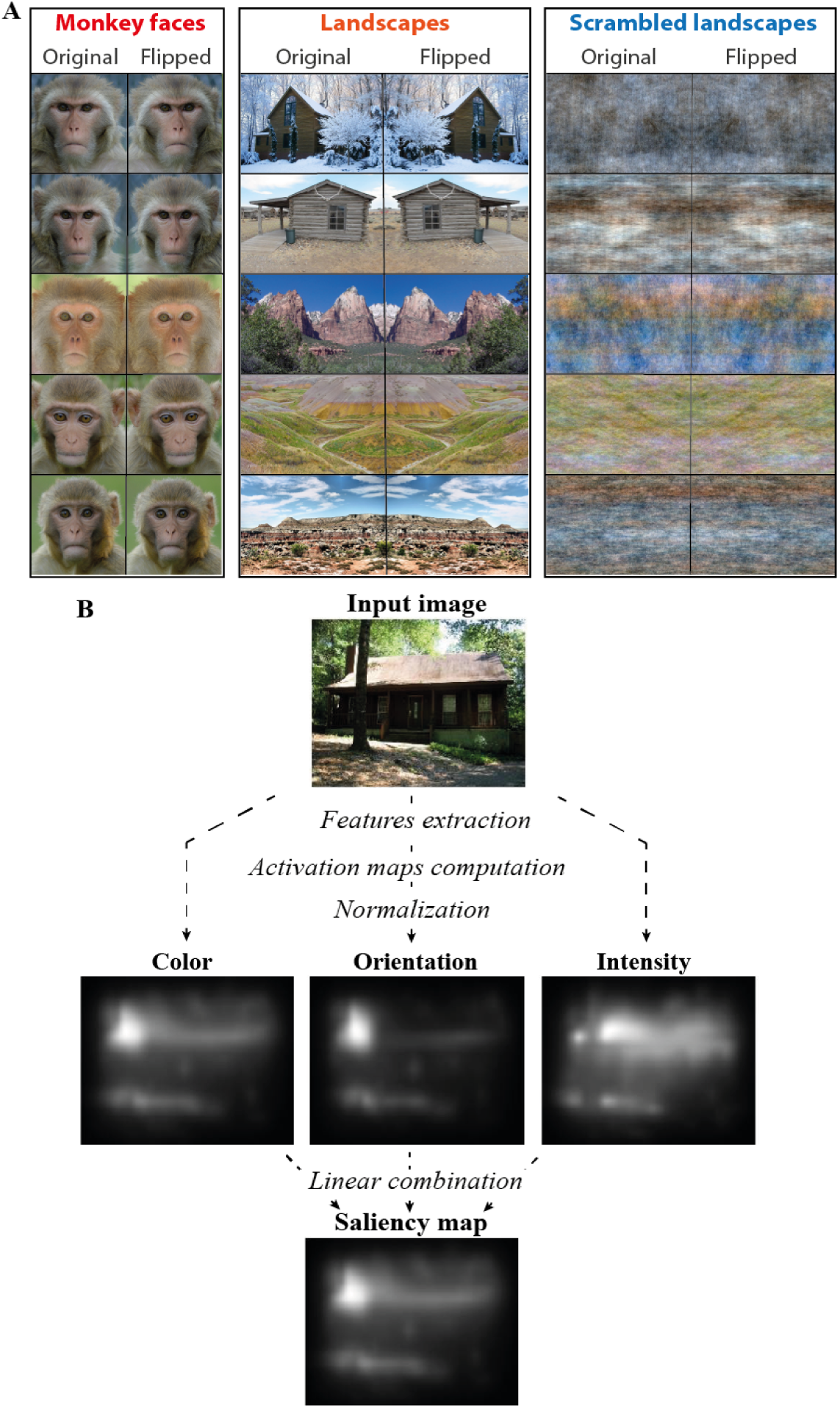
Image categories and GBVS model. **A**: Example of images presented during one testing session. Each testing session comprised the presentation of 30 images of 3 different categories: 10 monkey faces, 10 natural landscape images and 10 scrambled landscape images. Each category comprised 5 original images and 5 horizontally flipped images. **B**: To compute the image saliency map, we used the Graph-Based Visual Saliency model (GBVS, (Harel et al., 2006)). This model computes the activation maps based on several features (color, intensity and orientation), normalizes them, and finally combines all the maps into one single saliency map.

### 2.4. Drug administration

Once the animals were familiarized with the free viewing protocol and accustomed to intramuscular injections (using positive reinforcement training (Coleman et al., 2008)), atomoxetine (ATX, Tocris Bioscience, Ellisville, MO) and saline (control) administration sessions began. ATX is a potent NE reuptake inhibitor, as shown in previous studies (Bymaster et al., 2002; Koda et al., 2010). We chose the smallest efficient doses reported in our and others’ previous studies conducted in monkeys (Gamo et al., 2010; Carole Guedj et al., 2017b; Guedj et al., 2019; Reynaud et al., 2019). Each experiment started with one week of saline administration, followed by 2 or 4 weeks of testing with increasing doses of ATX (one dose per week): 0.5mg/kg and 1mg/kg (GE and CE) or 0.1mg/kg, 0.5mg/kg, 1mg/kg and 1.5mg/kg (CA and GU. ATX or saline was administered intramuscularly 30 min prior to testing. In total, for each animal, we collected 3 to 5 sessions with each dose of ATX and 3 to 5 sessions of saline condition.

### 2.5. Data analysis

The data were analyzed separately for each monkey. Eye movements were visually inspected with a customized toolbox implemented in MATLAB. Eye movements were recorded during the fixation and stimulus presentation period. To adjust the eye position with the center of the stimulus, we subtracted from each sample collected during the image presentation its baseline, i.e. the mean eye position during the fixation period.

#### 2.5.1. Pupil diameter

We computed the averaged normalized pupil diameter in the fixation period (500ms before the image onset), for each animal and each pharmacological condition. For each trial, the mean pupil diameter across this 500ms window was divided by the root mean square separately for each animal. These measures were then compared across pharmacological conditions.

#### 2.5.2. Explorations parameters

First, we calculated the total duration of the exploration as the percentage of time that monkeys spent exploring the image per trial, i.e. the number of samples recorded inside the stimulus-image (±2°) divided by the total number of samples recorded during image presentation. Then, we defined fixation events, using EyeMMV toolbox implemented in MATLAB (Krassanakis et al., 2014), as eye positions lasting at least 70ms within a location of 1.2° of radius. The first fixation was excluded from the analysis as it was a direct consequence of the preceding central fixation period. All fixations which fell outside the image were excluded from the analysis (±2° tolerance). We calculated the fixation number per trial and the duration of each fixation.

To assess the effects of ATX on the exploration parameters, we calculated for each trial within each category of images the difference between the mean fixation duration or number of fixations in ATX and saline conditions:

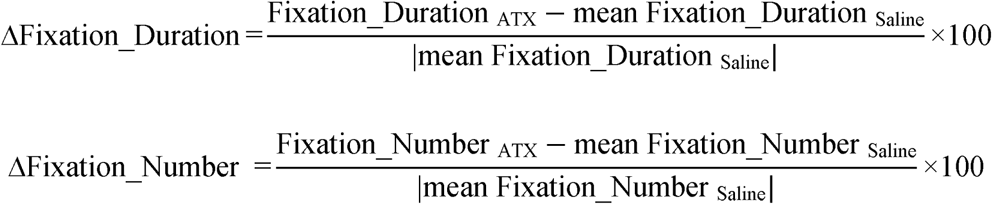

#### 2.5.3. Computation of the saliency-related fixations

We used the Graph-Based Visual Saliency model (GBVS) (Harel et al., 2006) to compute the saliency map of each image. The objective was to determine the impact of image salience on gaze orienting, which reflects the contribution of low-level sensory signals to attention control (Itti and Koch, 2000). This model uses a graph-based approach to obtain a saliency map that is dependent on global information. First, the feature maps are computed based on three dimensions (color, intensity and orientation). Then, activation maps are computed as a directional graph with edge weights depending on the dissimilarity or closeness of neighborhood nodes. Moreover, a distance penalty function is applied, so that nodes which are distant only weakly interact. These activation maps were then normalized to concentrate activation into a few key locations, and further combined into saliency maps (figure 1B). Based on these saliency maps, we calculated the mean saliency corresponding to each map and the mean saliency at each fixation (*saliency-related fixations*). We observed that the mean saliency was significantly different depending on the image category (*χ*^2^(_2_)=641.2, p<0.001). Specifically, the saliency was higher for the scrambled images compared to the intact images (landscape vs. scrambled: |t|_(27)_=16, p<0.001; scrambled vs. monkey face: |t|_(27)_=25, p<0.001), and the saliency was higher for the landscape images compared to the monkey face images (|t|_(27)_=9, p<0.001). For the *saliency-related fixations*, we calculated the mean value of saliency inside a square of 2° around the center of each fixation. The first fixation and all fixations outside the image were excluded from the analysis (±2° tolerance). Then, to assess the effects of ATX on the *saliency-related fixations* accounting for the image category bias in mean saliency, we normalized the *saliency-related fixations* by computing for each trial within each category of images the difference between the mean *saliency-related fixations* in ATX and saline conditions:

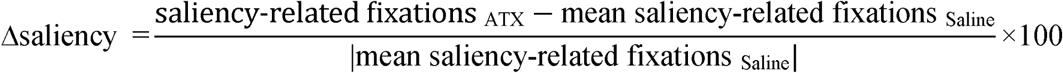

### 2.6. Statistical analysis

We used linear mixed models (using the ‘lme4’ package for R, (Bates et al., 2014)) to examine the effect of ATX on the different dependent measures described above, for each monkey. As a first step, we defined a model containing the most appropriate random effects (i.e. grouping factors and hierarchical structure of the data). Random effects were thus introduced sequentially, and their effect on model fit was assessed through Likelihood Ratio Tests (LRT): residuals of each model were compared, and the one with significantly lower deviance as assessed by a chi-squared test was chosen (Supplementary Table S1). We then tested the effect of the pharmacological condition and image category as fixed factors to evaluate the effect of ATX on the *exploration duration, fixation number* and *fixation duration*. Finally, post-hoc comparisons were carried out using pairwise comparisons through the ‘lsmeans’ package for R (Lenth, 2016), (p-adjusted with false discovery rate method (Benjamini and Hochberg, 1995)) to assess the effect of the different doses of ATX and the different categories of images.

Then, we used one-sample t-tests to determine whether the Δ*saliency* differed significantly from 0 in the ATX condition, i.e. to determine the influence of the different doses of ATX on *saliency-related fixations* with respect to the saline condition. To test the relationship between the *saliency-related fixations* in ATX and saline conditions, we used Spearman’s correlation tests including all monkeys to assess the correlation between the mean *saliency-related fixations* in saline condition and Δ*saliency*. Finally, we tested the relationship between Δ*saliency* and Δ*Fixation_duration* or Δ*Fixation_number* using a Spearman’s correlation test including all monkeys.

## 3. Results

### 3.1. Effect of ATX on pupil diameter (Figure 2)

To estimate the effect of ATX injection on LC-NE activity, we measured pupil diameter during the fixation period (500ms before the image onset). We found a significant main effect of pharmacological condition on pupil diameter in all monkeys (*χ*^2^_(4)_=697.9, p<0.001 for CA, *χ*^2^_(4)_=1004.3, p<0.001 for GU, *χ*^2^_(2)_=89.8, p<0.001 for CE, *χ*^2^_(2)_=46.1, p<0.001 for GE). All doses of ATX significantly increased pupil diameter compared to the saline condition (CA: |t|_(608.3)_=4.6, p<0.001 with 0.1mg/kg, |t|_(608.3)_=11.6, p<0.001 with 0.5mg/kg, |t|_(473.1)_=15.4, p<0.001 with 1mg/kg, |t|_(608.3)_=21.6, p<0.001 with 1.5mg/kg; GU: |t|_(711)_=4, p<0.001 with 0.1mg/kg, |t|_(711)_=13, p<0.001 with 0.5mg/kg, |t|_(712.3)_=18.2, p<0.001 with 1mg/kg |t|_(711)_=27.8, p<0.001 with 1.5mg/kg; CE: |t|_(443)_=2.8, p=0.005 with 0.5mg/kg, |t|_(437.4)_=9.2, p<0.001 with 1mg/kg; GE: |t|_(443)_=4.8, p<0.001 with 0.5mg/kg, |t|_(424.2)_=6.4, p<0.001 with 1mg/kg). Note that the effect size (computed as the difference between the mean pupil diameter in saline and ATX conditions) induced by ATX injection on pupil diameter differed between monkeys. Specifically, the increase of pupil diameter induced by ATX was lower for GE (maximum increase of 0.03 with 1mg/kg of ATX) compared to the other monkeys (1.0mg/kg of ATX: CA: 0.13, GU: 0.13, CE: 0.08).

**Figure 2.**
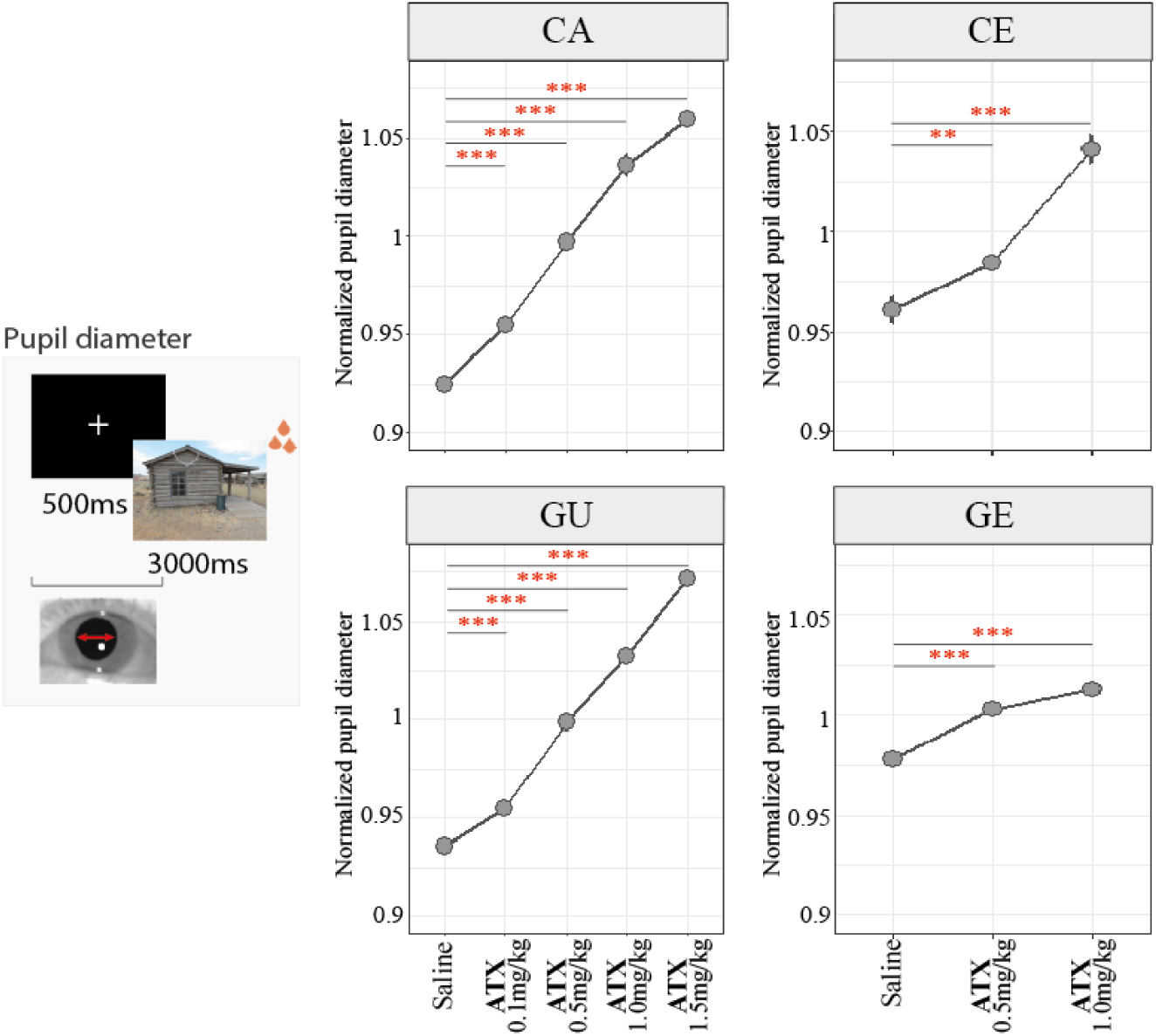
ATX effect on pupil diameter. For each animal and each pharmacological condition, we computed the averaged normalized (mean divided by the root mean square) pupil diameter (mean ± s.e.) during the fixation period (500ms before the image onset). ATX significantly increased pupil diameter as a function of the dose, in all monkeys, during the fixation period. **:p-value < 0.01; ***:p-value < 0.001.

### 3.2. Effect of ATX on the exploration parameters

To determine whether ATX injection modulates monkeys’ exploration behavior, we tested the effect of ATX on duration of exploration, duration of fixations and number of fixations. These results are summarized in Table 1 (see supplementary table S2 for more details). The most consistent effect of ATX across animals concerned the *fixation duration* (Figure 3). We found a significant main effect of pharmacological condition on *fixation duration* for 3 out of 4 monkeys (*χ*^2^_(4)_=10.7, p=0.03 for CA, *χ*^2^_(4)_=17.3, p=0.001 for GU and *χ*^2^_(2)_=21.5, p<0.001 for CE). Note that for monkey GE, with the lowest increase of pupil diameter after ATX injection, ATX did not impact fixation duration. ATX significantly increased the *fixation duration* in the other 3 monkeys (CA: |z|=4.1, p<0.001 with ATX 1mg/kg; GU: |z|=9.1, p<0.001 with ATX 0.5mg/kg, |z|=5.2, p<0.001 with ATX 1mg/kg, |z|=4.5, p<0.001 ATX 1.5mg/kg; CE: |t|_(2690)_=12.1, p<0.001 with ATX 0.5mg/kg). This effect was accompanied by a significant interaction between pharmacological condition and image category for two monkeys (*χ*^2^_(8)_=20.9, p=0.007 for GU and *χ*^2^_(4)_=61.9, p<0.001 for CE). Specifically, for monkey CE, *fixation duration* increased for monkey faces (t_(2694)_=-8.3, p<0.001) and scrambled landscape images (|t|_(2692)_=9.9, p<0.001) with ATX 0.5mg/kg whereas it decreased for landscapes (|t|_(2681)_=2.8, p=0.008) and scrambled landscape images (|t|_(2700)_=2.2, p=0.02) with ATX 1.0mg/kg. For monkey GU, *fixation duration* overall increased most consistently with ATX 0.5 mg/kg, in which the effect was detected for all image categories (|z|=7.4, p<0.001 for monkey face, |z|=3.4, p=0.003 for landscape and |z|=4.6, p<0.001 for scrambled landscape). Other doses were also associated with increased *fixation duration*, but less consistently across image categories (0.1mg/kg: |z|=2.1, p=0.04 for monkey face; 1mg/kg: z|=4.7, p<0.001 for monkey face; 1.5mg/kg: |z|=2.4, p=0.02 for monkey face, |z|=3.3, p=0.003 for scrambled landscape). In summary, the effect of the different doses of ATX on *fixation duration* varied across animals. For each animal, a given dose of ATX tended to enhance more effectively the *fixation duration* as compared to the other doses, without any systematic bias for a given category. For monkeys GU and CE, this effect was found with the ATX dose of 0.5mg/kg. For monkey CA, this effect was found with a higher dose of ATX, that is 1mg/kg.

**Table 1.** Effect of ATX on the exploration parameters. Results of pairwise comparisons between the saline and the doses of ATX with corrections for multiple comparisons (see supplementary table S2 for more details). ↗ or ↘: significant increase or decrease, respectively, after ATX administration; *: no significant interaction between pharmacological condition and image category; -: no significant main effect of pharmacological condition and no interaction between pharmacological condition and image category; NA: not applicable. Overall, ATX modulates all exploration parameters and the most consistent effect was found for the fixation duration.

|  |  | Total duration |  |  |  | Fixation number |  |  |  | Fixation duration |  |  |  |
| --- | --- | --- | --- | --- | --- | --- | --- | --- | --- | --- | --- | --- | --- |
|  |  | ATX01 | ATX05 | ATX10 | ATX15 | ATX01 | ATX05 | ATX10 | ATX15 | ATX01 | ATX05 | ATX10 | ATX15 |
| CA | Monkey face | - | - | - | - | - | - | - | - |  |  |  |  |
|  | Landscape | - | - | - | - | - | - | - | - | - | - | ↗* | - |
|  | Scrambled landscape | - | - | - | - | - | - | - | - |  |  |  |  |
| GU | Monkey face | - | - | - | - | - | - | - | - | ↗ | ↗ | ↗ | ↗ |
|  | Landscape | - | - | - | - | - | - | - | - | - | ↗ | - | ↗ |
|  | Scrambled landscape | - | - | - | - | - | - | - | - | - | ↗ | - | - |
| CE | Monkey face | NA | NA | ↗ | ↘ | NA | NA | ↘ | ↘ | NA | NA | ↗ | - |
|  | Landscape | NA | NA | - | ↘ | NA | NA | ↘ | - | NA | NA | - | ↘ |
|  | Scrambled landscape | NA | NA | ↗ | ↘ | NA | NA | ↘ | - | NA | NA | ↗ | ↘ |
| GE | Monkey face | NA | NA |  |  | NA | NA |  |  | NA | NA |  |  |
|  | Landscape | NA | NA | ↘* | ↗* | NA | NA | ↘* | ↗* | NA | NA | - | - |
|  | Scrambled landscape | NA | NA |  |  | NA | NA |  |  | NA | NA |  |  |

**Figure 3.**
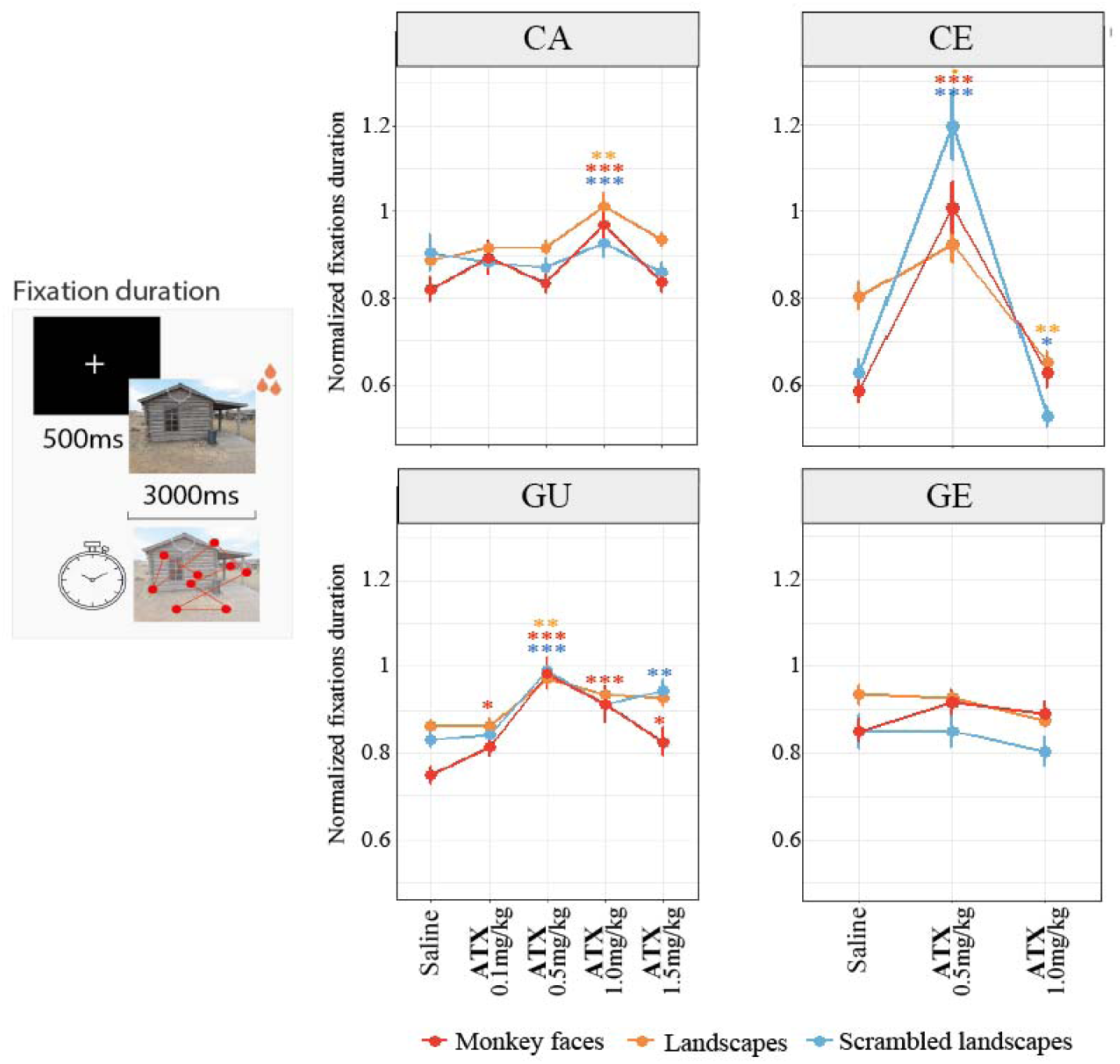
ATX effect on fixation duration. For each animal, each pharmacological condition and each image category, we computed the averaged normalized (mean divided by the root mean square) *fixation duration* (mean ± s.e.). Our results show that ATX increased the *fixation duration* for 3 out of 4 monkeys. *:p-value<0.05; **:p-value<0.01; ***:p-value<0.001.

The other exploration parameters, i.e. *exploration duration* and *fixation number*, were also impacted by ATX injections for two monkeys (CE and GE). Depending of the dose of ATX, the *exploration duration* was either increased (CE with 0.5mg/kg: |t|_(436)_=2.3, p=0.02 for scrambled landscape, |t|_(436)_=2.1, p=0.03 for monkey face; GE with 1mg/kg: |t|_(439.6)_=4.5, p<0.001 for all image categories) or decreased (CE with 1mg/kg: |t|_(439.7)_=4.2, p<0.001 for landscape, |t|_(439.7)_=3.4, p=0.001 for scrambled landscape, |t|_(439.7)_=4, p<0.001 for monkey face; GE with 0.5mg/kg: |t|_(436)_=2.7, p=0.006 for all image categories) compared to the saline condition. The *fixation number* decreased with 0.5mg/kg of ATX for monkeys CE (|t|_(431)_=2.8, p=0.01 for landscape, |t|_(431)_=6.1, p<0.001 for scrambled landscape and |t|_(431)_=5.4, p<0.001 for monkey face) and GE (|t|_(433)_=2.2, p=0.03 for all image categories), and with 1mg/kg for monkey face images for monkey CE (|t|_(433)_=3.3, p=0.002). For monkey GE, the *fixation number* increased with 1mg/kg of ATX (|t|_(437)_=3.9, p<0.001 for all image categories).

### 3.2. Effect of ATX on saliency-related fixations

Using the Graph-Based Visual Saliency model (Harel et al., 2006), we obtained the saliency map for each image, that reflects low-level features of the image in terms of three features (namely color, intensity and orientation). The saliency map is thus independent from monkeys’ behavior, rather depending on objective physical properties of the images. To investigate the impact of saliency on attentional orientating during free exploration, we computed the *saliency-related fixations*, corresponding to the mean value of saliency for each location the animals fixated.

Our results show that, in the saline condition, the animals’ gaze was differently guided by the saliency of the images, depending on the image category (Figure 4A). Specifically, the *saliency-related fixations* were higher for scrambled landscapes compared to intact landscapes and/or monkey face images for all monkeys (scrambled vs. intact landscape: |t|=3.1, p=0.002 for CA; |t|=5.2, p<0.001 for CE; |t|=7.3, p<0.001 for GE; scrambled landscape vs. face: |t|=7.3, p<0.001 for CA, |t|=5.3, p<0.001 for GU; |t|=10.1, p<0.001 for CE; |t|=8.1, p<0.001 for GE). The effect of ATX on *saliency-related fixations* was further assessed after normalizing the data (Δ*saliency*, see Methods section) to account for the image category bias (see methods) in mean saliency and *saliency-related fixations* in the saline condition. A Δ*saliency* above zero or below zero represents respectively, an increase or a decrease of *saliency-related fixations* following ATX injection. For scrambled images, the Δ*saliency* was significantly higher than zero for three monkeys with at least one dose of ATX (CA: |t|_(49)_=2.3, p=0.02 for 0.1mg/kg; GU: |t|_(49)_=3.5, p<0.001 for 0.1mg/kg; CE: |t|_(47)_=4.9, p<0.001 for 0.5mg/kg, |t|_(48)_=4.6, p<0.001 for 1mg/kg). The other monkey exhibited a decrease of Δ*saliency* for scrambled images (GE: |t|_(46)_=5.8, p<0.001 for 0.5mg/kg). On the contrary, ATX significantly decreased *saliency-related fixations* for intact images (landscapes and monkey faces) in two animals with at least one dose of ATX (CA: landscape: |t|_(49)_=2.9, p=0.006 for 1.5mg/kg; monkey face: |t|_(49)_=2.5, p=0.01 for 0.5mg/kg, |t|_(28)_=3.7, p<0.001 for 1mg/kg, |t|_(49)_=7.1, p<0.001 for 1.5mg/kg; GU: landscape: |t|_(39)_=2.3, p=0.02 for 1mg/kg; monkey face: |t|_(49)_=4.7, p<0.001 for 0.5mg/kg, |t|_(39)_=2.9, p=0.005 for 1mg/kg, |t|_(49)_=5.8, p<0.001 for 1.5mg/kg). The two other monkeys exhibited no effect of ATX for landscape images and an increase of *saliency-related fixations* for monkey face images (CE: |t|_(49)_=3.1, p=0.003 for 0.5mg/kg; GE: |t|_(49)_=2, p=0.048 for 0.5mg/kg). In summary, these results show that ATX modulated the animals’ exploration pattern based on the images category even after accounting for the difference of mean saliency between the three image-categories (Δ*saliency*) (Supplementary Table S3).

**Figure 4.**
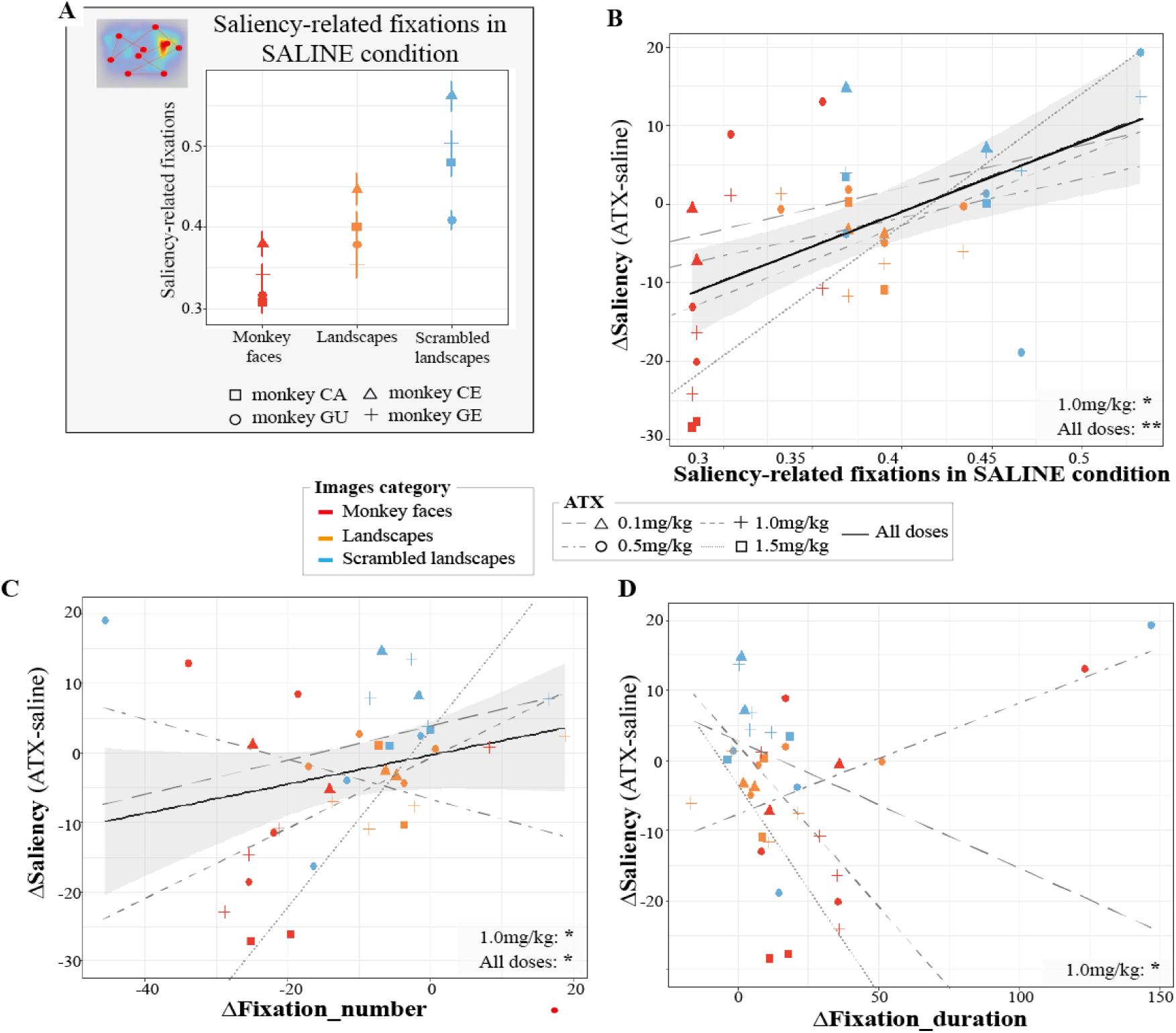
The effect of ATX on saliency-related fixations. **A:** For each monkey and each image category, we computed the *saliency-related fixations* in the saline condition. Our results show that the *saliency-related fixations* were higher for scrambled landscape compared to landscapeand/or monkey face images for all monkeys. **B:** For each animal, each pharmacological and each image category, we computed the Δsaliency as the difference between the *saliency-related fixations* in ATX and saline conditions. Our results show that ATX-induced changes in the *saliency-related fixations* was correlated with the *saliency-related fixations* in the saline condition with ATX 1.0mg/kg or combining all doses together (grey area, 95% CI combining all doses). **C:** For each animal, each pharmacological condition and each image category, we computed theΔnumber, as the difference between the fixations number in ATX and saline conditions. Our results show that ATX-induced changes in the *saliency-related fixations* was correlated with changes in the number of fixations with ATX 1.0mg/kg or combining all doses together (grey area, 95% CI combining all doses). **D:** For each animal, each pharmacological condition and each image category, we computed the Δduration, as the difference between the *fixation duration* in ATX and saline conditions. Our results show that ATX-induced changes in the *saliency-related fixations* was correlated with changes in the duration of fixations with ATX 1.0mg/kg. *:p-value<0.05; **:p-value<0.01.

To further characterize the effect of ATX, we examined the relationship between the *saliency-related fixations* in the saline condition and the Δ*saliency* for all doses of ATX. As illustrated in figure 4B, the difference in *saliency-related fixations* between ATX and saline conditions (Δ*saliency*) positively correlated with the mean *saliency-related fixations* in the saline condition (p=0.006, r=0.45). This result indicates that under ATX, the animals’ fixation pattern follows the same trend as that observed in the saline condition, with a stronger difference between images categories. ATX strengthened the exploratory pattern difference between image categories observed in the saline condition. In other words, ATX strengthen the contribution of saliency-driven gaze orienting when exploration in the control condition entails a high degree of saliency-driven orienting (scrambled landscape images) while ATX reduced the involvement of saliency-driven gaze orienting when exploration in the control condition entails a lower degree of saliency-driven gaze orienting (intact monkey face and landscape images).

Finally, to assess the link between the effect of ATX on exploration parameters and saliency-driven orienting, we examined the relationship between the Δ*saliency* and the Δ*fixation_number* or Δ*fixation_duration* for all doses of ATX. We found that the Δ*saliency* was positively correlated with the Δ*fixation_number* (p=0.02, r=0.32), i.e. the effect of ATX on the number of fixations. This relationship was present for all ATX doses and the slope of all these correlations tended to increase with the dose of ATX (but not with ATX 0.5 mg/kg where the pattern was reversed) (figure 4C). On the contrary, the Δ*saliency* was negatively correlated with the Δ*fixation_duration*, i.e. the effect of ATX on the duration of fixations, only for the highest common dose, i.e. 1.0mg/kg of ATX (p=0.014, r=-0.69) (figure 4D). In other words, under ATX, a larger number of fixations was associated with more *saliency-related fixations*, while longer fixations were associated with less *saliency-related fixation*s.

To sum up, ATX modulated the contribution of saliency-driven gaze orienting, quantified as the image-salience at each fixation (i.e. *saliency-related fixations)* depending on the image-category. Moreover, we found that the influence of ATX on saliency-driven gaze orienting correlated with the number and duration of fixations. Under ATX, depending on the image category, the animals either made more fixations of shorter duration to salient locations (in scrambled images), or less fixations with longer duration to salient locations (in intact monkey face and landscape images). Together, these results suggest that ATX modulated the contribution of low-level salience on attentional orienting as a function of the image category.

## 4. Discussion

We tested the impact of ATX, a NE reuptake inhibitor that increases NE availability in the brain, on attentional orienting during image exploration in four monkeys. First, we found that ATX impacted the way monkeys explored the images. The monkeys consistently spent more time on each fixation under ATX as compared to the control condition. Second, we found that ATX modulated the contribution of low-level signaling on spatial orienting, measured as the saliency-related fixations. Specifically, when exploration in the saline condition implies a high degree of saliency-driven orienting, i.e. for scrambled landscape images, ATX strengthen the saliency-driven orienting, while when exploration in the saline condition implies a low level of saliency-driven orienting, i.e. for monkey face and landscape images, ATX reduced the involvement of saliency-driven orienting. Moreover, this effect of ATX on saliency-driven orienting correlated with the effect of ATX on the number and duration of fixations. Our results suggest that NE adjusts the type of attentional orienting to the environment to explore.

### 4.1. Boosting NE transmission increases fixation duration regardless of the images content

We assessed the impact of ATX on the exploration pattern by measuring the total duration of exploration as well as the number and duration of fixations. We found that ATX consistently increases the fixation duration during the exploration of the different image categories (for 3 out of 4 monkeys). This effect varied as a function of the ATX dose and across individuals. This ATX dose-dependent effect was also found in the pupil diameter, with inter-individual variability. In summary, ATX increased pupil size for the three monkeys that showed an increase of duration of fixations while the pupil size was only slightly increased for the monkey that showed no effect of ATX injection on duration of fixations. While, we did not find any significant correlation between pupil size changes and fixation duration, the inter-individual variability induced by ATX might be related to differences in genetic determinants, in particular in a NE transporter gene (Greene et al., 2009; Hart et al., 2012; Kim et al., 2006; Whelan et al., 2012), and differences in neuronal and synaptic properties in response to neuromodulators (Hamood and Marder, 2014). Many studies showed that the duration of fixations varies with the image content (Rayner, 2009, 1998). For example, it has been shown that the duration of fixation was longer for atypical objects (Henderson et al., 1999) or when the scene luminance was decreased (Henderson et al., 2012; Loftus, 1985; Walshe and Nuthmann, 2014). When reading words, the fixation duration can be affected by different properties of words, such as their frequency, their predictability or their length (Kliegl et al., 2006; Reichle et al., 1998). The fixation duration reflects both visual and cognitive processing. Two possible interpretations can explain the increase of fixation duration after ATX injection. It can either reflect a slowdown of information processing, thus leading to longer fixations to process visual information, or it can reflect a deeper processing of visual information. On the basis of the facilitating effect of NE on sensory signal processing through an improvement of the signal-noise ratio in sensory cortex (C Guedj et al., 2017; Linster, 2019; Linster and Escanilla, 2019; Navarra and Waterhouse, 2019; Waterhouse and Navarra, 2019), we deem the first interpretation as unlikely. This role of the LC-NE system has been documented for the different sensory systems, visual (Navarra et al., 2013), auditory (Martins and Froemke, 2015), olfactory (Linster, 2019) and somatosensory system (Devilbiss and Waterhouse, 2004). Instead, the interpretation in terms of a deeper processing is more plausible. This increase in the duration of fixation was observed for all image categories tested, i.e. intact and scrambled landscapes and monkey faces, which leads us assuming an overall deeper processing of visual information regardless of the orienting process engaged during exploration. In other words, our results suggested that, in a free exploration context without any task requirement, ATX promotes a slower but more detailed visual processing of the environment (Glaholt and Reingold, 2012; Velichkovsky et al., 2000).

### 4.2. Boosting NE transmission adjusts the attentional orienting to the image content

During free exploration, overt attentional orienting inferred from gaze position can either be driven by low-level physical characteristics of the image or by higher-order signals such as the subjects’ preferences or interests, etc. To assess the degree of attentional capture induced by physical characteristics of the image, we computed the saliency map for each image based on their low-level features, i.e. color, intensity and orientation (Harel et al., 2006). Note that the animals viewed different intact and scrambled landscapes images every day, yet the same monkey face images were presented every day across all sessions. It is thus possible that familiarity with these latter images could differently impact the effect of ATX on gaze position. Despite this limitation, and as expected, in the saline condition we found that the animals’ gaze orientating was influenced by the level of saliency in the image [e.g. 50–52]. In the scrambled landscapes, with a higher mean saliency, the locations that the monkeys fixated were more salient compared to the intact images (landscape and monkey faces). We found that ATX strengthened this pattern. Specifically, for scrambled images, ATX increased the *saliency-related fixations* (for three monkeys) whereas for intact images (landscape and monkey faces), ATX decreased the *saliency-related fixations* (for two monkeys). In addition, a positive correlation was found between the effect of ATX on *saliency-related fixations* (Δ*saliency*) and the influence of saliency on gaze orienting in control condition (*saliency-related fixations* in the saline condition), which depends on the image category. In other word, ATX modulated the contribution of both types of orienting processes, i.e. saliency-driven and top-down-driven orienting, during image exploration. As discussed below, this finding fits with an increasing number of studies demonstrating the impact of NE onto sensory and high-level processes to shape behavior.

On the one hand, accumulating evidence has documented NE influences on sensory (bottom-up) processes, even at very early stages of sensory signal processing, improving the signal-noise ratio in sensory cortex in response to incoming stimuli of the environment (Navarra and Waterhouse, 2019; Waterhouse and Navarra, 2019). Recent studies showed that manipulating the NE level in humans modulates their perceptual sensitivity to detect a visual target (Gelbard-Sagiv et al., 2018; Guedj et al., 2019), and this effect reflected changes in evoked potentials and fMRI signals in the visual cortex (Gelbard-Sagiv et al., 2018). Another recent study showed that boosting the NE transmission in monkeys speeded up the target detection through a faster accumulation rate of sensory information (Reynaud et al., 2019). At rest, ATX was also found to reduce the functional correlation strength within sensory networks and to modify the functional connectivity between the LC and the fronto-parietal attention network (C Guedj et al., 2017; Carole Guedj et al., 2017a), involved in visuo-spatial orienting (Corbetta et al., 2008).

On the other hand, the LC-NE system is proposed to promote behavioral flexibility in order to adapt and optimize behavior depending on the contingencies of the environment (Aston-Jones et al., 1999; Aston-Jones and Cohen, 2005; Bouret and Sara, 2005), thus suggesting an impact of LC-NE system in high-level (top-down) processes. For example, the LC-NE system improves the ability to adapt behavior after a change in the rule in a set-shifting task in rats (Cain et al., 2011; McGaughy et al., 2008; Newman et al., 2008). A recent study also highlighted a context-dependent effect of the NE system, inferred from the pupil diameter, often considered as a proxy of the LC-NE activity: the authors reported larger diameter of the pupil in highly predictive contexts as compared to non-predictive contexts (Dragone et al., 2018). These context-dependent effects could result from the action of LC-NE system on prefrontal cortex that guides top-down driven behavior (Berridge and Spencer, 2016; Robbins and Arnsten, 2009).

Our results further reveal correlations between the effect of ATX on the *saliency-related fixations* and the effect of ATX on two exploration parameters, i.e. number and duration of fixations. Specifically, when ATX increased the *saliency-related fixations*, especially for scrambled images, it correlated with an increased number of fixations, and tended to increase fixation duration. By contrast, when ATX decreased the *saliency-related fixations*, especially for intact images, it correlated with a reduced number of fixations and longer fixation durations. This result suggests that ATX adjusts the attentional orienting to the image content. ATX strengthened the difference between the two types of attentional orienting observed in the saline condition: an automatic saliency-driven attention orienting, based on the physical properties of the images, and a more voluntary top-down-driven orienting, based on the subjects’ preferences and interests. These ATX-induced changes in attentional orienting could reflect a modulation of priority maps, a neural representation combining information of low-level saliency and top-down control. The LC-NE system could potentially act on the different brain areas involved in the computation of these priority maps (Bisley and Goldberg, 2010; Bisley and Mirpour, 2019; Fecteau and Munoz, 2006; Marsman et al., 2016; Mazer and Gallant, 2003; Mo et al., 2018) to bias information prioritization depending on the environment and adjust attentional orienting. As such, the LC-NE system would be in an ideal position to fine-tune behavior in order to appropriately and optimally respond to the environment and promote behavioral flexibility (Aston-Jones and Cohen, 2005; Navarra and Waterhouse, 2019).

In conclusion, our results reveal that, in naturalistic conditions, inhibiting the reuptake of norepinephrine with ATX injection adjusts the contribution of low-level salience on attentional orienting depending on the high-level image content. These results suggest that norepinephrine play a role in weighing the contribution of stimulus-driven and top-down control on attentional orienting.

## Acknowledgements

We thank Gislène Gardechaux and Frédéric Volland for technical and engineering assistance.

## Author Contributions

FHB conceived the work; FHB and AJR designed the work; AJR performed experiments and analyzed the data; EK, EB and EM provided support in data analysis; AJR and FHB interpreted the data and drafted the work; All authors revised and approved the final version of the manuscript.

## References

Arnsten, A.F.T., Wang, M.J., Paspalas, C.D., 2012. Neuromodulation of thought: flexibilities and vulnerabilities in prefrontal cortical network synapses. Neuron 76, 223–39. https://doi.org/10.1016/j.neuron.2012.08.038

Aston-Jones, G., Cohen, J.D., 2005. An integrative theory of locus coeruleus-norepinephrine function: adaptive gain and optimal performance. Annu. Rev. Neurosci. 28, 403–450. https://doi.org/10.1146/annurev.neuro.28.061604.135709

Aston-Jones, G., Rajkowski, J., Cohen, J., 1999. Role of locus coeruleus in attention and behavioral flexibility. Biol. Psychiatry 46, 1309–1320. https://doi.org/10.1016/S0006-3223(99)00140-7

Bates, D., Mächler, M., Bolker, B.M., Walker, S.C., 2014. Fitting linear mixed-effects models using lme4. 1406.5823v1.

Benjamini, Y., Hochberg, Y., 1995. Controlling the False Discovery Rate: A Practical and Powerful Approach to Multiple Testing. J. R. Stat. Soc. Ser. B. https://doi.org/10.2307/2346101

Berg, D.J., Boehnke, S.E., Marino, R.A., Munoz, D.P., Itti, L., 2009. Free viewing of dynamic stimuli by humans and monkeys 9, 1–15. https://doi.org/10.1167/9.5.19.Introduction

Berger, D., Pazienti, A., Flores, F.J., Nawrot, M.P., Maldonado, P.E., Gr??n, S., 2012. Viewing strategy of Cebus monkeys during free exploration of natural images. Brain Res. 1434, 34–46. https://doi.org/10.1016/j.brainres.2011.10.013

Berridge, C.W., Spencer, R.C., 2016. Differential cognitive actions of norepinephrine a2 and a1 receptor signaling in the prefrontal cortex. Brain Res. 1641, 189–196. https://doi.org/10.1016/j.brainres.2015.11.024

Bisley, J.W., Goldberg, M.E., 2010. Attention, intention, and priority in the parietal lobe. Annu. Rev. Neurosci. 33, 1–21. https://doi.org/10.1146/annurev-neuro-060909-152823

Bisley, J.W., Mirpour, K., 2019. The neural instantiation of a priority map. Curr. Opin. Psychol. 29, 108–112. https://doi.org/10.1016/J.COPSYC.2019.01.002

Bouret, S., Sara, S.J., 2005. Network reset: a simplified overarching theory of locus coeruleus noradrenaline function. Trends Neurosci. 28, 574–582. https://doi.org/10.1016/j.tins.2005.09.002

Bruce, N., Tsotsos, J., 2006. Saliency based on information maximization. Adv. Neural Inf. Process. Syst. 18 18, 155–162.

Buschman, T.J., Miller, E.K., 2007. Top-down versus bottom-up control of attention in the prefrontal and posterior parietal cortices. Science 315, 1860–1862. https://doi.org/10.1126/science.1138071

Bymaster, F.P., Katner, J.S., Nelson, D.L., Hemrick-Luecke, S.K., Threlkeld, P.G., Heiligenstein, J.H., Morin, S.M., Gehlert, D.R., Perry, K.W., 2002. Atomoxetine Increases Extracellular Levels of Norepinephrine and Dopamine in Prefrontal Cortex of Rat: A Potential Mechanism for Efficacy in Attention Deficit/Hyperactivity Disorder. Neuropsychopharmacology 27, 699–711. https://doi.org/10.1016/S0893-133X(02)00346-9

Cain, R.E., Wasserman, M.C., Waterhouse, B.D., McGaughy, J.A., 2011. Atomoxetine facilitates attentional set shifting in adolescent rats. Dev. Cogn. Neurosci. 1, 552–9. https://doi.org/10.1016/j.dcn.2011.04.003

Clark, C.R., Geffen, G.M., Geffen, L.B., 1989. Catecholamines and the covert orientation of attention in humans. Neuropsychologia 27, 131–139. https://doi.org/10.1016/0028-3932(89)90166-8

Coleman, K., Pranger, L., Maier, A., Lambeth, S.P., Perlman, J.E., Thiele, E., Schapiro, S.J., 2008. Training rhesus macaques for venipuncture using positive reinforcement techniques: a comparison with chimpanzees. J. Am. Assoc. Lab. Anim. Sci. 47, 37–41.

Corbetta, M., Patel, G., Shulman, G.L., 2008. The Reorienting System of the Human Brain: From Environment to Theory of Mind. Neuron 58, 306–324. https://doi.org/10.1016/j.neuron.2008.04.017

Corbetta, M., Shulman, G.L., 2002. Control of goal-directed and stimulus-driven attention in the brain. Nat. Rev. Neurosci. 3, 201–215. https://doi.org/10.1038/nrn755

Coull, J.T., Nobre, A.C., Frith, C.D., 2001. The Noradrenergic 2 Agonist Clonidine Modulates Behavioural and Neuroanatomical Correlates of Human Attentional Orienting and Alerting. Cereb. Cortex 11, 73–84. https://doi.org/10.1093/cercor/11.1.73

Devilbiss, D.M., Waterhouse, B.D., 2004. The Effects of Tonic Locus Ceruleus Output on Sensory-Evoked Responses ofVentral Posterior Medial Thalamic and Barrel Field Cortical Neurons in the Awake Rat. J. Neurosci. 24, 10773–10785. https://doi.org/10.1523/jneurosci.1573-04.2004

Dragone, A., Lasaponara, S., Pinto, M., Rotondaro, F., Luca, M. De, Doricchi, F., 2018. Expectancy modulates pupil size during endogenous orienting of spatial attention. CORTEX 102, 57–66. https://doi.org/10.1016/j.cortex.2017.09.011

Fecteau, J.H., Munoz, D.P., 2006. Salience, relevance, and firing□: a priority map for target selection 10. https://doi.org/10.1016/j.tics.2006.06.011

Gamo, N.J., Wang, M., Arnsten, A.F.T., 2010. Methylphenidate and Atomoxetine Enhance Prefrontal Function Through α2-Adrenergic and Dopamine D1 Receptors. J. Am. Acad. Child Adolesc. Psychiatry 49, 1011–1023. https://doi.org/10.1016/j.jaac.2010.06.015

Gelbard-Sagiv, H., Magidov, E., Sharon, H., Hendler, T., Nir, Y., 2018. Noradrenaline Modulates Visual Perception and Late Visually Evoked Activity. Curr. Biol. 28, 1–11. https://doi.org/10.1016/j.cub.2018.05.051

Glaholt, M.G., Reingold, E.M., 2012. Direct control of fixation times in scene viewing: Evidence from analysis of the distribution of first fixation duration. Vis. cogn. 20, 605–626. https://doi.org/10.1080/13506285.2012.666295

Greene, C.M., Bellgrove, M.A., Gill, M., Robertson, I.H., 2009. Noradrenergic genotype predicts lapses in sustained attention. Neuropsychologia 47, 591–594. https://doi.org/10.1016/j.neuropsychologia.2008.10.003

Guedj, Carole, Meunier, D., Meunier, M., Hadj-Bouziane, F., 2017a. Could LC-NE-Dependent Adjustment of Neural Gain Drive Functional Brain Network Reorganization? Neural Plast. 2017, 1–12. https://doi.org/10.1155/2017/4328015

Guedj, Carole, Monfardini, E., Reynaud, A.J., Farnè, A., Meunier, M., Hadj-bouziane, F., 2017b. Boosting Norepinephrine Transmission Triggers Flexible Recon figuration of Brain Networks at Rest 4691–4700. https://doi.org/10.1093/cercor/bhw262

Guedj, C, Monfardini, E., Reynaud, A.J., Farnè, A., Meunier, M., Hadj-Bouziane, F., 2017. Boosting Norepinephrine Transmission Triggers Flexible Reconfiguration of Brain Networks at Rest. Cereb. Cortex 27, 4691–4700. https://doi.org/10.1093/cercor/bhw262

Guedj, C., Reynaud, A., Monfardini, E., Salemme, R., Farnè, A., Meunier, M., Hadj-Bouziane, F., 2019. Atomoxetine modulates the relationship between perceptual abilities and response bias. Psychopharmacology (Berl). https://doi.org/https://doi.org/10.1007/s00213-019-05336-7

Hadj-Bouziane, F., Monfardini, E., Guedj, C., Gardechaux, G., Hynaux, C., Farnè, A., Meunier, M., 2014. The helmet head restraint system: A viable solution for resting state fMRI in awake monkeys. Neuroimage 86, 536–543. https://doi.org/10.1016/j.neuroimage.2013.09.068

Hamood, A.W., Marder, E., 2014. Animal-to-Animal Variability in Neuromodulation and Circuit Function. Cold Spring Harb. Symp. Quant. Biol. 79, 21–8. https://doi.org/10.1101/sqb.2014.79.024828

Harel, J., Koch, C., Perona, P., 2006. Graph-Based Visual Saliency. Neural Inf. Process. Syst. 19, 545:552.

Hart, A.B., de Wit, H., Palmer, A.A., 2012. Genetic factors modulating the response to stimulant drugs in humans. Curr. Top. Behav. Neurosci. 12, 537–77. https://doi.org/10.1007/7854_2011_187

Henderson, J.M., 2017. Gaze Control as Prediction. Trends Cogn. Sci. https://doi.org/10.1016/j.tics.2016.11.003

Henderson, J.M., Nuthmann, A., Luke, S.G., 2012. Eye Movement Control During Scene Viewing: Immediate Effects of Scene Luminance on Fixation Durations. https://doi.org/10.1037/a0031224

Henderson, J.M., Weeks, P.A., Hollingworth, A., 1999. The effects of semantic consistency on eye movements during complex scene viewing. J. Exp. Psychol. Hum. Percept. Perform. 25, 210–228. https://doi.org/10.1037/0096-1523.25.1.210

Itti, L., Baldi, P., 2005. A principled approach to detecting surprising events in video. Proc. - 2005 IEEE Comput. Soc. Conf. Comput. Vis. Pattern Recognition, CVPR 2005 I, 631–637. https://doi.org/10.1109/CVPR.2005.40

Itti, L., Koch, C., 2000. A saliency-based search mechanism for overt and covert shifts of visual attention. Vision Res. 40, 1489–1506.

Itti, L., Koch, C., Niebur, E., 1998. A Model of Saliency-Based Visual Attention for Rapic Scene Analysis. IEEE Trans. Pattern Anal. Mach. Intell. 20, 1254–1259. https://doi.org/10.1109/34.730558

Kim, C.-H., Hahn, M.K., Joung, Y., Anderson, S.L., Steele, A.H., Mazei-Robinson, M.S., Gizer, I., Teicher, M.H., Cohen, B.M., Robertson, D., Waldman, I.D., Blakely, R.D., Kim, K.-S., 2006. A polymorphism in the norepinephrine transporter gene alters promoter activity and is associated with attention-deficit hyperactivity disorder. Proc. Natl. Acad. Sci. U. S. A. 103, 19164–9. https://doi.org/10.1073/pnas.0510836103

Kliegl, R., Nuthmann, A., Engbert, R., 2006. Tracking the mind during reading: The influence of past, present, and future words on fixation durations. J. Exp. Psychol. Gen. 135, 12–35. https://doi.org/10.1037/0096-3445.135.1.12

Koda, K., Ago, Y., Cong, Y., Kita, Y., Takuma, K., Matsuda, T., 2010. Effects of acute and chronic administration of atomoxetine and methylphenidate on extracellular levels of noradrenaline, dopamine and serotonin in the prefrontal cortex and striatum of mice. J. Neurochem. 114, no-no. https://doi.org/10.1111/j.1471-4159.2010.06750.x

Krassanakis, V., Filippakopoulou, V., Nakos, B., 2014. EyeMMV toolbox□: An eye movement post-analysis tool based on a two-step spatial dispersion threshold for fixation identification. J. Eye Mov. Res. 7, 1–10. https://doi.org/10.16910/jemr.7.1.1

Lenth, R. V., 2016. Least-Squares Means: The R Package lsmeans [WWW Document].

Linster, C., 2019. Cellular and network processes of noradrenergic modulation in the olfactory system. Brain Res. 1709, 28–32. https://doi.org/10.1016/j.brainres.2018.03.008

Linster, C., Escanilla, O., 2019. Noradrenergic effects on olfactory perception and learning. Brain Res. 1709, 33–38. https://doi.org/10.1016/j.brainres.2018.03.021

Loftus, G.R., 1985. Picture Perception: Effects of Luminance on Available Information and Information-Extraction Rate. Psychological Association, Inc.

Marsman, J.C., Cornelissen, F.W., Dorr, M., Renken, R.J., 2016. A novel measure to determine viewing priority and its neural correlates in the human brain. J. Vis. 16, 1–18. https://doi.org/10.1167/16.6.3.doi

Martins, A.R.O., Froemke, R.C., 2015. Coordinated forms of noradrenergic plasticity in the locus coeruleus and primary auditory cortex. Nat. Neurosci. 18, 1483. https://doi.org/10.1038/NN.4090

Mazer, J.A., Gallant, J.L., 2003. Goal-related activity in V4 during free viewing visual search. Evidence for a ventral stream visual salience map. Neuron 40, 1241–50. https://doi.org/10.1016/s0896-6273(03)00764-5

McGaughy, J., Ross, R.S., Eichenbaum, H., 2008. Noradrenergic, but not cholinergic, deafferentation of prefrontal cortex impairs attentional set-shifting. Neuroscience 153, 63–71. https://doi.org/10.1016/j.neuroscience.2008.01.064

Mo, C., He, D., Fang, F., 2018. Attention Priority Map of Face Images in Human Early Visual Cortex. J. Neurosci. 38, 149–157. https://doi.org/10.1523/JNEUROSCI.1206-17.2017

Navarra, R.L., Clark, B.D., Zitnik, G.A., Waterhouse, B.D., 2013. Methylphenidate and atomoxetine enhance sensory-evoked neuronal activity in the visual thalamus of male rats. Exp. Clin. Psychopharmacol. 21, 363–374. https://doi.org/10.1037/a0033563.Methylphenidate

Navarra, R.L., Waterhouse, B.D., 2019. Considering noradrenergically mediated facilitation of sensory signal processing as a component of psychostimulant-induced performance enhancement. Brain Res. https://doi.org/10.1016/j.brainres.2018.06.027

Newman, L.A., Darling, J., McGaughy, J., 2008. Atomoxetine reverses attentional deficits produced by noradrenergic deafferentation of medial prefrontal cortex. Psychopharmacology (Berl). 200, 39–50. https://doi.org/10.1007/s00213-008-1097-8

Oliva, A., Torralba, A., 2007. The role of context in object recognition. Trends Cogn. Sci. https://doi.org/10.1016/j.tics.2007.09.009

Parkhurst, D., Law, K., Niebur, E., 2002a. Modeling the role of salience in the allocation of overt visual attention. Vision Res. 42, 107–123. https://doi.org/10.1016/S0042-6989(01)00250-4

Parkhurst, D., Law, K., Niebur, E., 2002b. Modeling the role of salience in the allocation of overt visual attention. Vision Res. 42, 107–123. https://doi.org/10.1016/S0042-6989(01)00250-4

Rayner, K., 2009. Eye movements and attention in reading, scene perception, and visual search, Quarterly Journal of Experimental Psychology. https://doi.org/10.1080/17470210902816461

Rayner, K., 1998. Psychological Bulletin Eye Movements in Reading and Information Processing: 20 Years of Research. Psychological Association, Inc.

Reichle, E.D., Pollatsek, A., Fisher, D.L., Rayner, K., 1998. Toward a Model of Eye Movement Control in Reading. Psychol. Rev. 105, 125–157. https://doi.org/10.1037/0033-295X.105.1.125

Reynaud, A.J., Froesel, M., Guedj, C., Ben Hadj Hassen, S., Cléry, J., Meunier, M., Ben Hamed, S., Hadj-Bouziane, F., 2019. Atomoxetine improves attentional orienting in a predictive context. Neuropharmacology 150, 59–69. https://doi.org/10.1016/J.NEUROPHARM.2019.03.012

Robbins, T.W., Arnsten, A.F.T., 2009. The Neuropsychopharmacology of Fronto-Executive Function: Monoaminergic Modulation. Annu. Rev. Neurosci. 267–287. https://doi.org/10.1146/annurev.neuro.051508.135535.The

Velichkovsky, B.M., Dornhoefer, S.M., Pannasch, S., Unema, P.J.A., 2000. Visual fixations and level of attentional processing. Proc. Eye Track. Res. Appl. Symp. 2000 79–85. https://doi.org/10.1145/355017.355029

Walshe, C.R., Nuthmann, A., 2014. Asymmetrical control of fixation durations in scene viewing. Vision Res. 100, 38–46. https://doi.org/10.1016/j.visres.2014.03.012

Waterhouse, B.D., Navarra, R.L., 2019. The locus coeruleus-norepinephrine system and sensory signal processing□: A historical review and current perspectives. Brain Res. 1–15. https://doi.org/10.1016/j.brainres.2018.08.032

Whelan, R., Conrod, P.J., Poline, J.-B., Lourdusamy, A., Banaschewski, T., Barker, G.J., Bellgrove, M.A., Büchel, C., Byrne, M., Cummins, T.D.R., Fauth-Bühler, M., Flor, H., Gallinat, J., Heinz, A., Ittermann, B., Mann, K., Martinot, J.-L., Lalor, E.C., Lathrop, M., Loth, E., Nees, F., Paus, T., Rietschel, M., Smolka, M.N., Spanagel, R., Stephens, D.N., Struve, M., Thyreau, B., Vollstaedt-Klein, S., Robbins, T.W., Schumann, G., Garavan, H., 2012. Adolescent impulsivity phenotypes characterized by distinct brain networks. Nat. Neurosci. 15, 920–925. https://doi.org/10.1038/nn.3092

Zhou, B., Lapedriza, A., Khosla, A., Oliva, A., Torralba, A., 2017. Places□: A 10 million Image Database for Scene Recognition 8828, 1–14. https://doi.org/10.1109/TPAMI.2017.2723009

Zhou, B., Lapedriza, A., Xiao, J., Torralba, A., Oliva, A., 2014. Learning deep features for scene recognition using places database. Adv. Neural … 487–495. https://doi.org/10.1162/153244303322533223

