## Supplementary information for "Atomoxetine modulates the contribution of high- and low-level signals during free viewing of natural images in rhesus monkeys"

### Supplementary table S1

**Linear mixed models**. For each variable, we defined a model containing the most appropriate random effects (i.e. grouping factors and their possible nesting) and we tested the effects of the fixed factors (i.e. experimental manipulations). Random effects were sequentially introduced, and their effect on model fit was assessed through Likelihood Ratio Tests (see methods).

| Variables | Family of linear mixed model | Random factors | Fixed factors |
|---|---|---|---|
| **Duration of exploration** | Gaussian | Session and mirror image | Pharmacological condition and image category |
| **Number of fixations** | Gaussian | Session and mirror image | Pharmacological condition and image category |
| **Duration of fixations** | Gaussian | Session, mirror image and trial | Pharmacological condition and image category |
| **Saliency-related fixations** | Gaussian | Session, mirror image and trial | Pharmacological condition and image category |
| **Normalized pupil diameter** | Gaussian | Session | Pharmacological condition |
| **Saliency map** | Gaussian | maps | Image category |

### Supplementary table S2

**Effect of ATX on the exploration parameters.** P-values reflect pairwise comparisons between the saline and the doses of ATX with corrections for multiple comparisons. ↗ or ↘: significant increase or decrease, respectively, after ATX administration; *: no significant interaction between pharmacological condition and image category, p-values reflect main effect of pharmacological condition; -: no significant main effect of pharmacological condition and no interaction between pharmacological condition and image category.

**Total duration**

| | | **saline** | **ATX01** | | | **ATX05** | | | **ATX10** | | | **ATX15** | | |
|---|---|---|---|---|---|---|---|---|---|---|---|---|---|---|
| | | Mean | Mean | $p_{fdr}$ | effect | Mean | $p_{fdr}$ | effect | Mean | $p_{fdr}$ | effect | Mean | $p_{fdr}$ | effect |
| **CA** | **Monkey face** | 87.7 | 73.6 | - | - | 73.2 | - | - | 75.5 | - | - | 71.7 | - | - |
| | **Landscape** | 96.8 | 94.2 | - | - | 95.2 | - | - | 94.6 | - | - | 97 | - | - |
| | **Scrambled landscape** | 81.4 | 78 | - | - | 77.9 | - | - | 75.3 | - | - | 74.6 | - | - |
| **GU** | **Monkey face** | 68.6 | 66.4 | - | - | 66.9 | - | - | 63.3 | - | - | 62.8 | - | - |
| | **Landscape** | 86.5 | 83.1 | - | - | 86 | - | - | 86.2 | - | - | 86.4 | - | - |
| | **Scrambled landscape** | 68.8 | 65.7 | - | - | 69.7 | - | - | 74.2 | - | - | 75.9 | - | - |

| | | **saline** | **ATX05** | | | **ATX10** | | |
|---|---|---|---|---|---|---|---|---|
| | | Mean | Mean | $p_{fdr}$ | effect | Mean | $p_{fdr}$ | effect |
| **CE** | **Monkey face** | 92.7 | 98.3 | **0.03** | ↗ | 82.4 | **<0.001** | ↘ |
| | **Landscape** | 98 | 93.4 | 0.08 | - | 86.8 | **<0.001** | ↘ |
| | **Scrambled landscape** | 89.9 | 95.8 | **0.02** | ↗ | 81.1 | **0.001** | ↘ |
| **GE** | **Monkey face** | 72.7 | 65.5 | | | 82.9 | | |
| | **Landscape** | 81.5 | 80.6 | **0.006*** | ↘ | 88.8 | **<0.001*** | ↗ |
| | **Scrambled landscape** | 78.8 | 67 | | | 88.9 | | |

**Fixation number**

| | | **saline** | **ATX01** | | | **ATX05** | | | **ATX10** | | | **ATX15** | | |
|---|---|---|---|---|---|---|---|---|---|---|---|---|---|---|
| | | Mean | Mean | $p_{fdr}$ | effect | Mean | $p_{fdr}$ | effect | Mean | $p_{fdr}$ | effect | Mean | $p_{fdr}$ | effect |
| **CA** | **Monkey face** | 8.1 | 6.1 | - | - | 6.3 | - | - | 5.7 | - | - | 6.1 | - | - |
| | **Landscape** | 11 | 10.4 | - | - | 10.6 | - | - | 9.5 | - | - | 10.6 | - | - |
| | **Scrambled landscape** | 6.4 | 6.3 | - | - | 6.3 | - | - | 5.8 | - | - | 6 | - | - |
| **GU** | **Monkey face** | 7.2 | 6.2 | - | - | 5.4 | - | - | 5.4 | - | - | 5.8 | - | - |
| | **Landscape** | 10.4 | 9.8 | - | - | 9.4 | - | - | 9.5 | - | - | 9.7 | - | - |
| | **Scrambled landscape** | 7.3 | 6.8 | - | - | 6.4 | - | - | 7.3 | - | - | 7.3 | - | - |

| | | **saline** | **ATX05** | | | **ATX10** | | |
|---|---|---|---|---|---|---|---|---|
| | | Mean | Mean | $p_{fdr}$ | effect | Mean | $p_{fdr}$ | effect |
| **CE** | **Monkey face** | 7.2 | 4.8 | **<0.001** | ↘ | 5.7 | **0.002** | ↘ |
| | **Landscape** | 7.7 | 6.4 | **0.012** | ↘ | 7.5 | 0.73 | - |
| | **Scrambled landscape** | 6.1 | 3.3 | **<0.001** | ↘ | 5.9 | 0.71 | - |
| **GE** | **Monkey face** | 5.8 | 4.7 | | | 6.3 | | |
| | **Landscape** | 8 | 8 | **0.029*** | ↘ | 9.5 | **<0.001*** | ↗ |
| | **Scrambled landscape** | 4.6 | 3.8 | | | 5.3 | | |

| Fixation duration | | | | | | | | | | | | | | |
|---|---|---|---|---|---|---|---|---|---|---|---|---|---|---|
| | | **saline** | **ATX01** | | | **ATX05** | | | **ATX10** | | | **ATX15** | | |
| | | Mean | Mean | $p_{fdr}$ | effect | Mean | $p_{fdr}$ | effect | Mean | $p_{fdr}$ | effect | Mean | $p_{fdr}$ | effect |
| **CA** | **Monkey face** | 269.5 | 293.8 | | | 274.7 | | | 319.3 | | | 275.2 | | |
| | **Landscape** | 213.8 | 220.5 | 0.25* | - | 220.9 | 0.89* | - | 244.1 | **<0.001*** | ↗ | 225.3 | 0.89* | - |
| | **Scrambled landscape** | 298.1 | 290.7 | | | 286.5 | | | 305.7 | | | 283 | | |
| **GU** | **Monkey face** | 220.4 | 240.2 | **0.04** | ↗ | 290.1 | **<0.001** | ↗ | 268.9 | **<0.001** | ↗ | 243.6 | **0.02** | ↗ |
| | **Landscape** | 194.2 | 194.2 | 0.95 | - | 218.9 | **0.003** | ↗ | 210.7 | 0.08 | - | 208.6 | 0.08 | ↗ |
| | **Scrambled landscape** | 207.6 | 210.6 | 0.74 | - | 248.1 | **<0.001** | ↗ | 227.9 | 0.06 | - | 235.9 | **0.003** | - |

| | | **saline** | **ATX05** | | | **ATX10** | | |
|---|---|---|---|---|---|---|---|---|
| | | Mean | Mean | $p_{fdr}$ | effect | Mean | $p_{fdr}$ | effect |
| **CE** | **Monkey face** | 293.8 | 505.1 | **<0.001** | ↗ | 314.3 | 0.66 | - |
| | **Landscape** | 294.1 | 337.5 | 0.05 | - | 239.1 | **0.007** | ↘ |
| | **Scrambled landscape** | 319.4 | 610.3 | **<0.001** | ↗ | 268.4 | **0.02** | ↘ |
| **GE** | **Monkey face** | 311.2 | 335.1 | | | 326.2 | | |
| | **Landscape** | 244.7 | 242.8 | - | - | 229.5 | - | - |
| | **Scrambled landscape** | 407.4 | 408.5 | | | 384.8 | | |

### Supplementary table S3

**Effect of ATX on Δsaliency.** P-values from one-sample t-tests to determine whether the Δsaliency (ATX vs. saline), for the different doses of ATX, differed significantly from 0.

| ΔSaliency (ATX vs. saline) | | | | | | | |
|---|---|---|---|---|---|---|---|
| | | **Monkey face** | | **Landscape** | | **Scrambled landscape** | |
| | | mean ΔSaliency | p-value | mean ΔSaliency | p-value | mean ΔSaliency | p-value |
| **CA** | 0.1mg/kg | -0.56 | 0.91 | -3.9 | 0.36 | **7.1** | **0.02** |
| | 0.5mg/kg | **-13.1** | **0.01** | -4.9 | 0.22 | 1.4 | 0.71 |
| | 1.0mg/kg | **-24.1** | **<0.001** | -7.5 | 0.16 | 6.8 | 0.07 |
| | 1.5mg/kg | **-28.4** | **<0.001** | **-10.9** | **0.006** | 0.1 | 0.98 |
| **GU** | 0.1mg/kg | -7.2 | 0.12 | -3.4 | 0.53 | **14.7** | **<0.001** |
| | 0.5mg/kg | **-20.1** | **<0.001** | 1.9 | 0.64 | -3.8 | 0.33 |
| | 1.0mg/kg | **-16.4** | **0.005** | **-11.6** | **0.02** | 3.9 | 0.39 |
| | 1.5mg/kg | **-27.7** | **<0.001** | 0.3 | 0.93 | 3.4 | 0.32 |
| **CE** | 0.5mg/kg | **13** | **0.002** | -0.2 | 0.95 | 19.3 | **<0.001** |
| | 1.0mg/kg | -10.8 | 0.08 | -6.1 | 0.25 | 13.7 | **<0.001** |
| **GE** | 0.5mg/kg | **8.9** | **0.048** | -0.6 | 0.88 | **-18.9** | **<0.001** |
| | 1.0mg/kg | 1.2 | 0.75 | 1.3 | 0.79 | 4.3 | 0.18 |
